# Interpretable Drug Target Predictions using Self-Expressiveness

**DOI:** 10.1101/2021.03.01.433365

**Authors:** Diego Galeano, Santiago Noto, Ruben Jimenez, Alberto Paccanaro

**Affiliations:** School of Applied Mathematics, Fundação Getulio Vargas, Rio de Janeiro, Brazil; Department of Computer Science, Royal Holloway, University of London, Egham, UK

## Abstract

The identification of missing drug targets is critical for the development of treatments and for the molecular elucidation of drug side effects. Drug targets have been predicted by exploiting molecular, biological or pharmacological features of drugs and protein targets. Yet, developing integrative and interpretable machine learning models for predicting drug targets remains a challenging task. We present Inception, an integrative and interpretable matrix completion model for predicting drug targets. Inception is a self-expressive model that learns two similarity matrices: one for drugs and another for protein targets. These learned similarity matrices are key for our models’ interpretability: they can explain how a predicted drug-target interaction can be explain in terms of a linear combination of chemical, biological and pharmacological similarities. We develop a novel objective function with efficient closed-form solution. To demonstrate the ability of Inception at recovering missing drug-target interactions (DTIs), we perform cross-validation experiments with stringent controls of data imbalance, chemical similarities between drugs and sequence similarities between targets. We also assess the performance of our model using a simulated prospective approach. Having trained our model with DTIs from a snapshot 2011 of the DrugBank database, we test whether we could predict DTIs from a 2020 snapshot of DrugBank. Inception outperforms two state-of-the-art drug target prediction models in all the scenarios. This suggests that Inception could be useful for predicting missing drug target interactions while providing interpretable predictions.

## 1 Introduction

The identification of drug targets is critical for the elucidation of drugs therapeutic effects, the anticipation of drug side effects, and even for the prevention of drug resistance (Schenone et al., 2013). Typically, a limited number of drug targets are experimentally identified through costly and time consuming *in-vitro* experiments. Yet, our knowledge of drug targets remains largely incomplete [Yildirim et al., 2007].

A wide range of computational approaches have been proposed to predict missing drug-target interactions (DTI). Early work focuses on docking simulations and ligand-based approaches. Docking simulations attempt to model physico-chemical drug-protein interactions by considering the 3D structure of the proteins. However, these simulations are computationally expensive, often requiring expert knowledge, and they can only be used when the 3D protein structure is available (see review in [Amaro et al., 2018]). Ligand-based methods (Keiser et al., 2007) relate protein targets based on the chemical similarity of their ligands and then use statistical models to make predictions. Yet, these methods often neglect other relevant information, such as protein sequence.

Machine learning approaches that have been proposed to predict missing drug-targets can be divided into two main groups. A first group of methods are based on exploiting the structure of the drug-target network, in combination with chemical structure similarities between drugs and protein sequence similarities between targets [Yamanishi et al., 2008, Bleakley and Yamanishi, 2009, Mei et al., 2013, Chen et al., 2012]. The drug-target network is a bipartite graph where nodes represent drugs and targets and edges represent experimentally known interactions. For instance, Bleakley and Yamanishi, 2009 use a support vector machine classifier with kernels based on drug chemical similarities and target sequence similarities. Chen et al., 2012 build a heterogeneous network by adding further links to the drug-target network: weighted links between pairs of drugs representing their chemical and shared targets similarities, and weighted links between pairs of targets representing their sequences and shared drugs similarities. In the heterogeneous network, starting from a given drug, they perform a random walk with restart to rank the candidate targets for the given drug. The limitation of these methods is that they only integrate chemical information about drugs and sequence information about their protein targets. However, other type of information, such as drug side effects, have also been useful in the prediction of missing drug targets [Campillos et al., 2008].

The second group of machine learning methods integrate a wider range of heterogeneous information about drugs and their protein targets. In an early attempt, Zheng et al., 2013 propose a collaborative matrix factorisation model that uses drug similarities based on their chemical structures and Anatomical, Therapeutic and Chemical (ATC) classification, and protein similarities based on their similarities in protein sequence, gene ontology annotations and protein interactions. Recently, Luo et al., 2017 proposed DTINet, a network integration model that exploits seven types of heterogeneous data about drugs and targets, including chemical structure, drug-drug interactions, drug side effects, disease indications, protein sequence, protein-protein interactions and protein-disease associations. DTINet has two main learning steps. In the first, it learns a low-dimensional vector representation for drugs and targets based on the heterogeneous information using diffusion component analysis [Cho et al., 2015]. In the second step, the learned representations are used as features in an inductive matrix completion model, where interaction is predicted based on the geometric closeness between the drugs and the targets. Wan et al., 2019 proposed neoDTI, that improves on DTINet by using a neural network model to learn topology-preserving embeddings of drugs and targets from the heterogeneous network.

Building integrative machine learning models it is not an straightforward task, and it remains an open challenge across biology and medicine [Zitnik et al., 2019]. DTINet and neoDTI have effectively combined a wide range of heterogeneous information about drugs and targets to provide state-of-the-art predictions. Yet, even these accurate models were not designed for providing interpretable predictions; which is critical to aid decision making in drug discovery [Rudin, 2019, Jiménez-Luna et al., 2020].

Here we present Inception, an integrative machine learning model that predicts missing DTIs more accurately than previously proposed prediction models. Inception is a matrix completion model inspired by recent high-rank matrix completion under self-expressive models (SEM) [Elhamifar, 2016]. The core idea of SEM models is to represent datapoints (e.g. drugs) as a linear combination of few other datapoints. This is different from the representation used by DTINet and neoDTI where drugs and targets are represented by learned low-dimensional feature vectors. Importantly, we constrain our matrix completion to learn from non-negative similarities between drugs and targets. This has the advantage of offering interpretable predictions in terms of available chemical, biological and pharmacological information. Our results indicate that Inception provides state-of-the-predictions and that it might be useful to provide human-in-the-loop interpretations of the predictions.

## 2 Methods

Self-expressive models are based on the *self-expressiveness* property of the data [Elhamifar and Vidal, 2013], which states that each data point in the union of subspaces can be efficiently represented as a linear combination of other data points [Fan and Chow, 2017, Frasca et al., 2019, Galeano and Paccanaro, 2019]. Given an incomplete data matrix 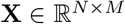, SEM models aims to represent each column of **X** as a sparse representation of other columns, i.e. **X** ≃ **XW**, where 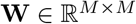 denotes the self-representation coefficient matrix for the column elements. A similar model can be obtained for the rows of **X**, i.e. **X** ≃ **HX**, where 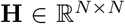 denotes the self-representation coefficient matrix for the row elements. Typically, diag(**W**) = 0 (or diag(**W**) = 0) is enforced to prevent the trivial solution of representing a datapoint by itself.

To learn the self-representation matrices **W** and/or **H**, previous work focused on solving sparse optimization programs under *l_1_* relaxation. Galeano and Paccanaro [2019], Frasca et al. [2019] also proposed Geometric SEM (GSEM), a framework to integrate side information using graph regularisation constraints. Yet, these models do not effectively integrate multiple side information about **X** and their learned self-representation matrices, **W** and/or **H**, are difficult to interpret because they are not symmetric, i.e. similarities.

We propose Inception, an integrative SEM model that constrains the self-representation matrices to learn from similarities relating drugs and targets. Inception exploits the use of well-known similarity information between drugs, such as chemical structure similarity, and between protein targets, such as sequence similarity, to provide interpretable predictions in terms of known available chemical, biological or pharmacological information. Our starting point is the matrix **X** containing binary drug-target interactions between *N* drugs (rows) and *M* protein targets (columns). An entry *x_ij_* = 1 if drug *i* was found to experimentally bind to protein target *j*, or *x_ij_* = 0 otherwise. Our task consist on predicting the presence or absence of drug-target interactions in **X**. Inception predicts scores by weighting the contributions of the SEM model for drugs and targets, as follow:

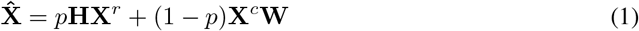

where **X**^*r*^ and **X**^*c*^ are obtained after normalising rows and columns of **X** such that the norm of each row (or column) is one, respectively; and *p* ∈ [0,1] controls for the relative importance of each model for the prediction.

Figure 1 illustrates Inception’s model in Equation (1). The figure depicts (in blue) how our model integrates several drug similarities to learn **H**, including information from chemical structure (denoted as 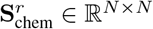), drug indication 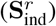, drug side effects 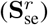 and drug-drug interactions 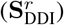. An additional similarity matrix is built based on the cosine similarity between the row elements of the normalised DTI matrix 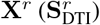. Similarly, Figure 1 shows (in red) how our model integrates several target similarities to learn **W**, including information from protein sequence 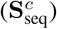, protein-protein interactions 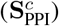, and protein-disease association 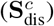. A similarity matrix based on the cosine similarity between the columns of the normalised DTI matrix 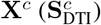 is also build.

**Figure 1:**
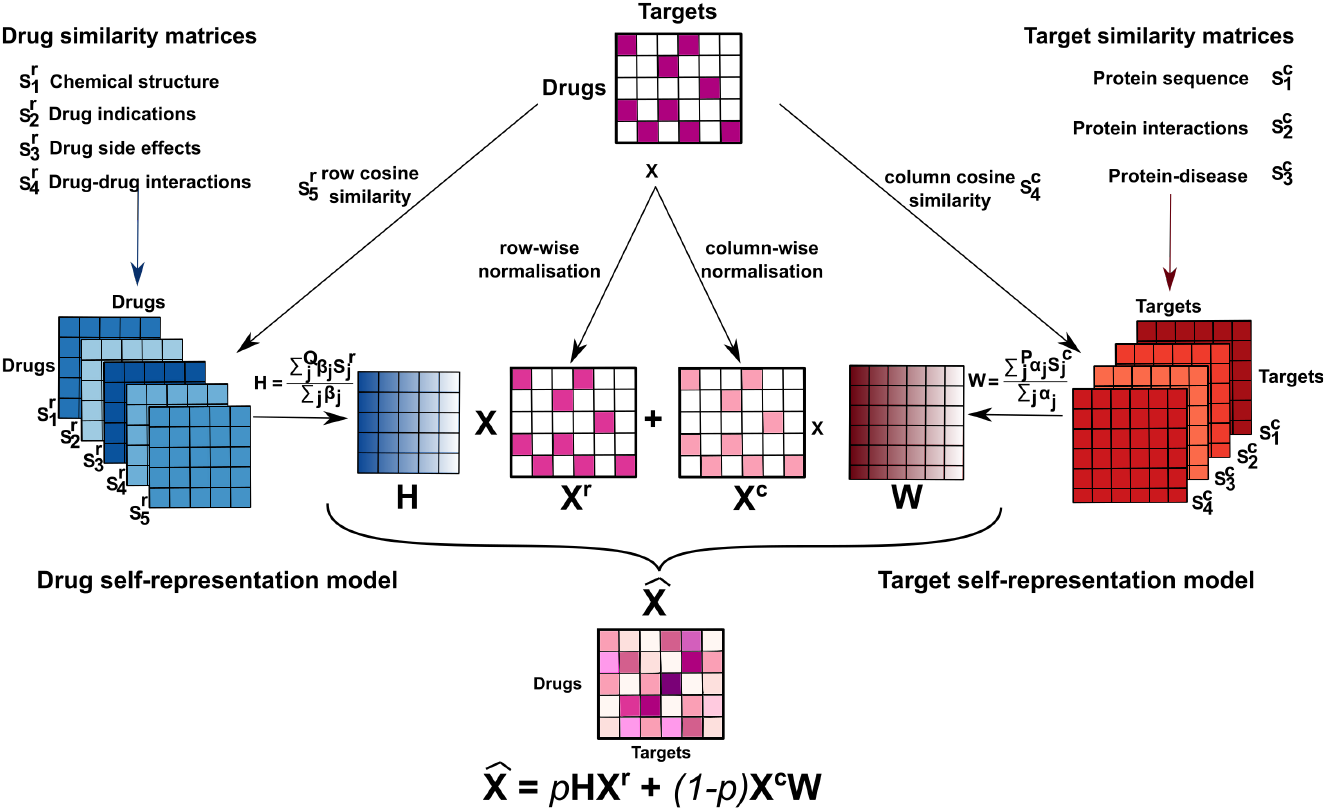
Overview of our approach. 1, 923 drug-target interactions (DTIs) between 549 drugs and 424 protein targets were used. The DTIs were arranged into an *n x m* matrix **X** by encoding them with values of 1 (pink). Unobserved associations were encoded with zeroes (white). Our algorithm, Inception, is the linear combination of two self-expressive models. The drug self-representation model, **HX**^*r*^, represents each drug (row of **X**), as a linear combination of other drugs by learning a similarity matrix **H**, that linearly aggregates similarity matrices relating drugs (shown in blue). The target self-representation model, **X**^*c*^ **W**, represents each target (column of **X**), as a linear combination of other targets by learning a similarity matrix **W**, that linearly aggregates similarity matrices relating protein targets (shown in red). Inception’s predicts scores by combining both linear models 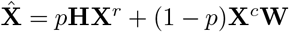, where *p* is a constant value that controls the importance of each model for the prediction.

Inception learns **W** and **H** from Equation 1 by minimising the following cost function:

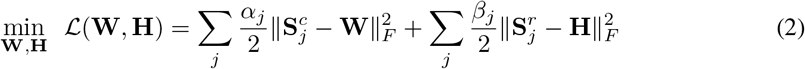

where ||·||_F_ denotes the Frobenius norm, and *α_j_ >* 0 and *β_j_ >* 0 are constant values. The first term in Equation 2 is the target self-representation constraint, that ensures that **W** is a weighted linear combination of similarity matrices relating protein targets. Notice how the constant values *α_j_* weigh the importance of each similarity matrix for **W**. This informs us about the relative importance of each type of side information for the prediction. Similarly, the second term in Equation 2, the drug self-representation constraint, ensures that **H** learns from the weighted linear combination of similarity matrices relating drugs. The objective function in Equation 2 is convex in both **W** and **H**. This allows us to obtain a closed form solution for our predictions 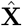. By setting 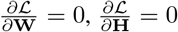, we obtain the optimal solution for **W** and **H**:

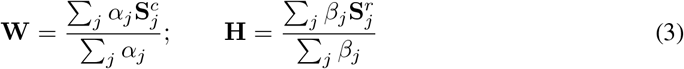

By replacing Equation (3) in (1), our predicted scores 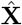 can be written as:

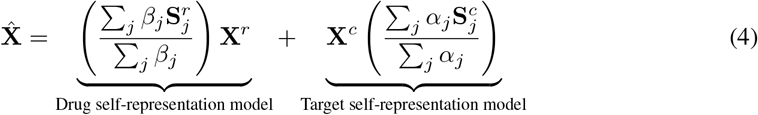

Thus our approach does not require iterations of any optimisation procedure, it can be computed extremely fast, and it scales up to large drug-target interaction datasets with multiple heterogeneous sources of information.

## 3 Materials

### Datasets

We use the datasets compiled by Luo et al., 2017 in their original study. It includes Drug-Target Interactions (DTIs) from a 2011 snapshot of DrugBank v3.0 (Knox et al., 2010) and complementary information about drugs and protein targets. It contains 1923 DTIs between 549 drugs and 424 protein targets. Each drug and each protein in this set have at least one known association. Complementary information about drugs includes 2D Tanimoto chemical similarity, drug-drug interactions from DrugBank v3.0 (Knox et al., 2010), drug-disease associations from the Comparative Toxicogenomics database (Davis et al., 2013) and drug-side effects from the Side Effect Resource (SIDER) v2.0 (Kuhn et al., 2010). Complementary information about proteins includes sequence similarity, protein-protein interactions from the Human Protein Reference Database (HPRD) (Keshava Prasad et al., 2009), and protein-disease association from the Comparative Toxicogenomics database (Davis et al., 2013). In total, there are seven heterogeneous sources of complementary information. Details about each dataset can be found in Supplementary Table 1.

### Similarity matrices

Similarities between drugs are obtained as follows. 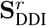 is obtained after computing the cosine similarity between the row elements of the 549 x 549 drug-drug interaction (DDI) matrix. Similarly, we obtain 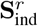 and 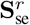 from the 549 x 5,022 drug-disease association matrix and the 549 x 3, 588 drug-side effect association matrix, respectively. We use the same 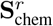 provided by Luo et al., 2017, which consists on the 2D Tanimoto chemical similarity. 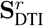 is also computed based on the cosine similarity between the rows of *X^r^*. In total, we obtain five similarity matrices relating drugs based on diverse chemical, biological and pharmacological data.

To obtain similarities between protein targets, we also compute the cosine similarity between protein target features based on their 424 x 5, 022 protein-disease association matrix, and 424 x 424 protein-protein interaction data; from which we build 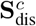 and 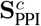, respectively. The sequence similarity matrix 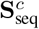, which is based on the Smith-Waterman percent identity score provided by Luo et al., 2017, is normalised between 0 and 1. 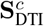 is also computed based on the cosine similarity between the columns of *X^c^*. In total, we obtain four similarity matrices relating proteins based on genomic and proteomic features.

The similarity values in each of the nine similarity matrices, 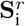 and 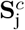, are bounded between 0 and 1. This helps in the statistical interpretation of the contributions of each individual type of information to the optimal *W* and *H*, by inspecting the values of *α_s_* and *β_s_* in Equation (2). While high similarity values can be informative of the biology underlying a drug-target interaction, low similarity values might represent noisy information that can be removed. To achieve a sparser and more meaningful solution for interpretability, we define thresholds values *τ*_chem_, *τ*_se_, *τ*_ind_, *τ*_seq_, *τ*_dis_ for the 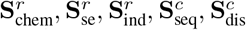 similarity matrices, respectively. Below each threshold, the values in the similarity matrices were set to zero. These values are defined in the following section.

### Evaluation procedure

We compare the performance of Inception with two state-of-the-art integrative DTI prediction models: DTINet [Luo et al., 2017] and neoDTI [Wan et al., 2019]. Following their procedure, we frame a binary classification problem, where the goal is to predict the presence or absence of drug-target interactions. We perform a ten-fold cross validation procedure where 10% of the known DTIs in *X* are randomly chosen as a test set. The remaining 90% of the DTIs are used for training. Notice that the values of *X* that are removed for testing are set to a value of zero in the DTI matrix used for training. The prediction performance is then assessed based on the ability of the model to discriminate interacting pairs versus non-interacting pairs in the test set. We use the area under the receiving operator curve (AUROC) and the area under the precision recall curve (AUPR) to measure the binary classification performance.

To set the model parameters, we perform a grid search on a validation set in each fold. On average, we observe good results with *α*_seq_ = 5 (*τ*_seq_ = 0.245), *α*_ppi_ = 2, *α*_dis_ = 5 (*τ*_dis_ = 0.99), *β*_chem_ = 1 (*τ*_chem_ = 0.684), *β*_se_ = 3.5 (*τ*_se_ = 0.574), *β*_ind_ = 9.5 (*τ*_ind_ = 0.866), *β*_chem_ = 0.5. Note that we do not tune *α*_DTI_ = 1 and *β*_DTI_ = 1, that correspond to the penalisation values for 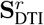 and 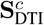, respectively. We use these hyperparameters for all our experiments.

To guarantee a fair comparison, we tune the model parameters of each model using the same folds that are used for our method. In detail,

- **DTINet**: we use the code provided by Luo et al., 2017. We perform a grid search of hyperparameters for DTINet and find an optimal performance when setting the dimension of drugs and proteins embeddings to 150, and the restart probability of the random walk to 0.5. For the matrix completion, we set the rank to 40 and the regularisation parameter to 10.
- **NeoDTI**: we use the code provided by Wan et al., 2019, which we updated to the latest version of Tensorflow 2.1 [Abadi et al., 2015]. Following the guidelines provided by Wan et al., 2019, we set the dimension of the embeddings and projection matrices to 1024, which gives the best performance for our splits.

## 4 Results

### Performance evaluation on multiple drugs

Following Luo et al., 2017, we start by analysing the performance of Inception at recovering missing drug-target associations in **X** in a balanced scenario. During the ten fold cross-validation, we sample at random a matching number of non-interacting pairs (negative labels) with respect to the number of interacting pairs (positive labels) in the test set. Results show that Inception significantly outperforms state-of-the-art competitors. Inception achieves an average AUROC of 95.96 ± 1.00% (mean and s.t.d.), which represents a performance gain of 6.5% with respect to neoDTI (89.43 ± 2.22%) and DTINet (89.79 ± 1.85%). The increase in prediction performance is also significantly higher in terms of AUPR, for which our method achieves 96.25 ± 0.73%, versus 90.16 ± 01.94% and 91.57 ± 1.24% for neoDTI and DTINet, respectively. These results suggest that our method is able to effectively recover missing DTIs in **X**.

In practice, there are more non-interacting pairs in **X** than interacting pairs (0.83% of the associations in **X** are interacting pairs). To simulate the more realistic scenario where interacting pairs are recovered from a larger sample of non-interacting pairs, we expanded the evaluation presented by Luo et al., 2017 and evaluate the performance of the methods at predicting missing DTIs by increasing the ratio of negative to positive labels. We randomly sample the set of non-interacting pairs in the test set such that the ratios of negative to positive labels are 1, 5, 10, 20, 50 and 100. Figure 2a shows that the mean AUROC of the three methods is robust with respect to the class imbalance. Inception outperforms neoDTI and DTINet across all the ratios. In terms of AUPR, which is known to be sensitive to class imbalance [Saito and Rehmsmeier, 2015], we observe that the performance of the three methods decrease around 10% when the ratio of negative to positive labels is increased by 10 (see Fig. 2b). We also observe that DTINet is more sensitive to the class imbalance than Inception or neoDTI. Exact AUROC and AUPR values are shown in Supplementary Tables 2-3.

**Figure 2:**
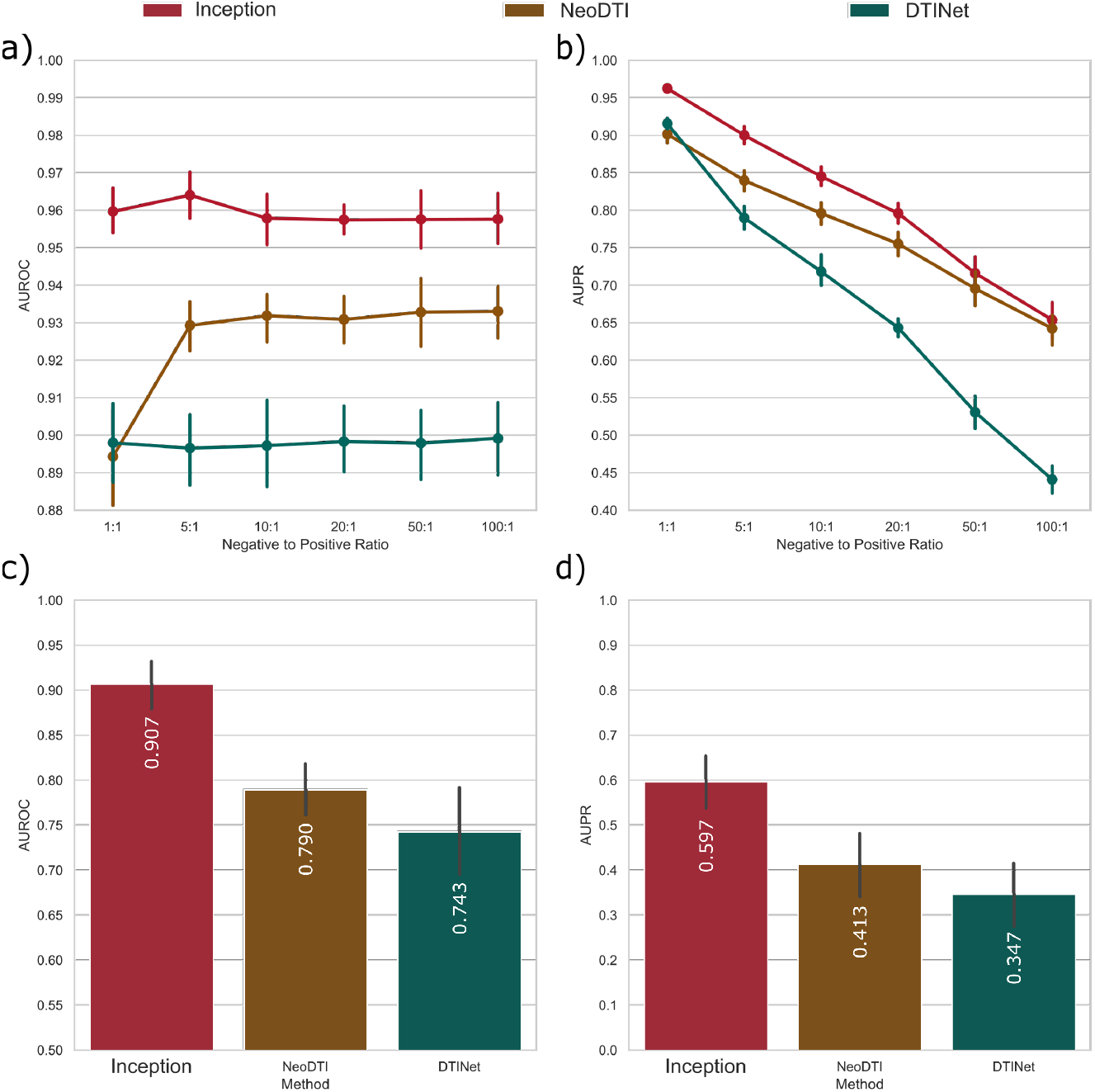
Evaluation of our model at predicting missing drug-target interactions in stringent scenarios. The performance of Inception (our method) is compared to the state-of-the-art drug target prediction models DTINet and neoDTI. **(a)** mean area under the receiver operating curve (AUROC) performance (*y*-axis) of each method when controlling for data imbalance in the test set. The negative to positive ratio (*x*-axis) was obtained by a negative sampling of the unknown DTIs in the test set. Error bars indicate the standard deviation. **(b)** mean area under the precision-recall (AUPR) curve performance (*y*-axis) of each method when controlling for data imbalance in the test set. **(c-d)** mean AUROC and AUPR performance when predicting drug target interactions for chemically dissimilar drugs (Tanimoto chemical similarity below 0.6) and protein targets with dissimilar sequence (sequence identity below 40%). Error bars indicate the standard deviation.

An interesting question is whether chemically similar drugs could bias our evaluations of the method’s performance. This may occur because the distribution of chemical similarities for drugs that share targets is significantly higher than for those that do not (One-Tailed Wilcoxon Sum Rank Significance, *p <* 2.23 x 10^−308^). Following Luo et al., 2017, we assess the performance of the methods using the same test sets from our cross-validation sets, for a class imbalance with ratio 10, but where the DTIs corresponding to drugs with more than 0.6 chemical similarity to drugs in the training set are removed. Our results show that Inception can predict more accurately missing DTIs in **X** even for drugs that are chemically dissimilar to the drugs available for training. The mean AUROC only decreases by 2.15% for Inception, while this percentage is 4.44% for neoDTI and 7.05% for DTINet. The performance drops in terms of AUPR are consistent: 8.57% (Inception), 10.77% (neoDTI) and 16.36% (DTINet). Exact AUROC and AUPR values are shown in Supplementary Tables 4-5.

Another interesting question is whether targets with similar sequence could bias our evaluations of the method’s performance. This is also motivated by the observation that the distribution of sequence similarities for proteins that share drugs is significantly higher than for those that do not (One-Tailed Wilcoxon Sum Rank Significance, *p <* 2.23 x 10 ^−308^). Following Luo et al., 2017, we remove DTIs in the test set corresponding to protein targets with sequence similarity higher than 40% by keeping a class imbalance with ratio 10. Our results show that Inception can predict more accurately missing DTIs in **X** even for targets that are dissimilar in sequence to the targets available for training. The mean AUROC only decreases by 3.9% for Inception, while this percentage is 8.57% for neoDTI and 10.4% for DTINet. The performance drops in terms of AUPR are also also consistent: 20.93% (Inception), 27.72% (neoDTI) and 29.7% (DTINet). Exact AUROC and AUPR values are shown in Supplementary Tables 4-5.

Following Luo et al., 2017, we also investigated the method’s performance when simultaneously controlling for chemically similarity and sequence similarity. We remove DTIs in the test set corresponding to protein targets with sequence similarity higher than 40% and drugs with chemical similarity higher than 0.6, and keeping a class imbalance with ratio 10. While the prediction performance of Inception only drops by 5.11% in terms of mean AUROC, the decline in performance is 14.23% for neoDTI and 15.44% for DTINet. The result is consistent in terms of mean AUPR: 24.85% (Inception), 38.31% (neoDTI) and 37.14% (DTINet). AUROC and AUPR values from the cross-validation procedure are shown in Figure 2c-d. Exact AUROC and AUPR values are shown in Supplementary Tables 4-5.

### Prospective Evaluation

We have shown that Inception effectively predicts missing DTIs in **X** under several evaluation procedures. Cross-validation procedures can be overoptimistic, as they do not mimic the experimental drug discovery process [Cami et al., 2011]. In practice, DTIs are continuously added to the DrugBank database by pharmacological experts as these are discovered and experimentally validated by the scientific community.

Using DTI data from DrugBank 2011, we build **X**, trained the models, and aim to predict novel DTIs that appear in DrugBank 2020 but were not known in 2011. This amounts to a prospective evaluation that preserves the chronological order in which information become historically available. We found 471 new DTIs between 169 drugs and 129 protein targets that were placed in a test set. Following Luo et al., 2017, we calculate the performance using the recall of novel DTIs found in the top-N predictions retrieved for each drug.

Figure 3 shows that, in 70% of the cases, Inception predicts the correct DTIs in the top-40 predictions retrieved. The percentage of DTIs in the top-40 predictions retrieved is significantly lower for neoDTI and DTINet. This means that many of these novel DTIs, which were discovered experimentally, could have been systematically predicted by our approach already in 2011.

**Figure 3:**
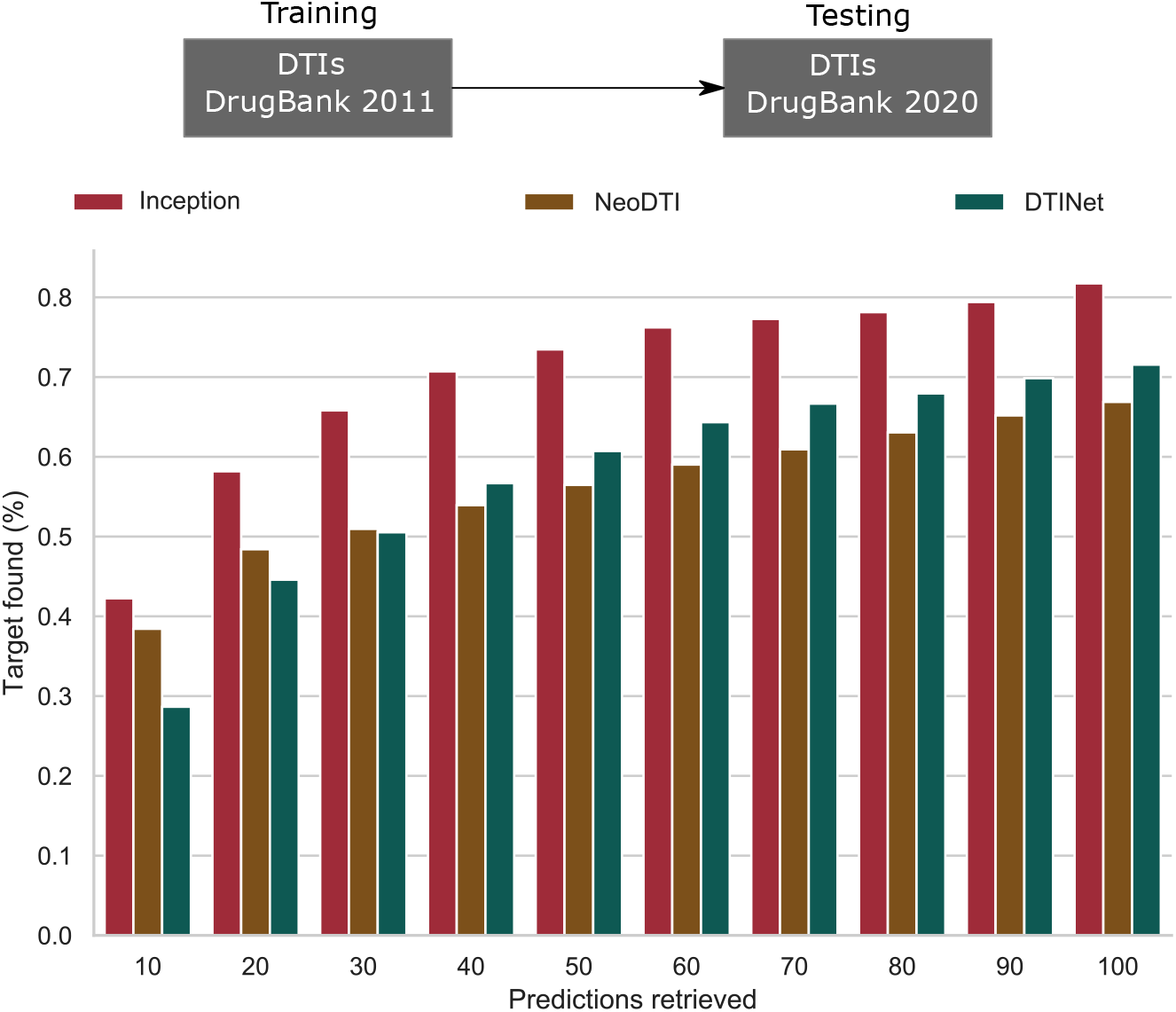
Prospective performance evaluation. Each model was trained with drug-target interactions (DTIs) from a 2011 snapshot of the DrugBank database to predict missing DTIs that were found in the 2020 snapshot. 471 novel DTIs between 169 drugs and 129 protein targets from the 2020 snapshot were used as test set. For each drug, we check whether the method was able to retrieved the correct target in the top-10, 20, 30, 40, 50, 60, 70, 80, 90 or 100 proteins retrieved. Our method outperform the competitors in all the cases. Inception (red), NeoDTI (brown) and DTINet (green).

### Interpreting Inception’s predictions

The effectiveness of Inception at predicting missing DTIs prompted us to ask whether our model can provide biologically meaningful interpretation for the predictions.

Let us recall that Inception is a linear model where the predicted scores are given by the linear combination of two terms (see Equation 1): the drug self-representation model (*HX^r^*) and the target self-representation model (*X^c^ W*). We started by analysing the contribution of each of these terms for a single DTI that was missing in our 2011 snapshot but appeared in the 2020 snapshot. We select the beta-blocker Atenolol (DB00335), for which we had only one known target in our 2011 snapshot: the beta-1 adrenergic receptor (ADRB1). In the 2020 snapshot, Atenolol was also associated to the beta-2 adrenergic receptor (ADRB2), its new target. The pair Atenolol-ADRB2 is in fact Inception’s top-1 prediction with a score of 1.141. 95% of this score is coming from the drug self-representation model, and only 5% from the target self-representation model (see Figure 4a).

**Figure 4:**
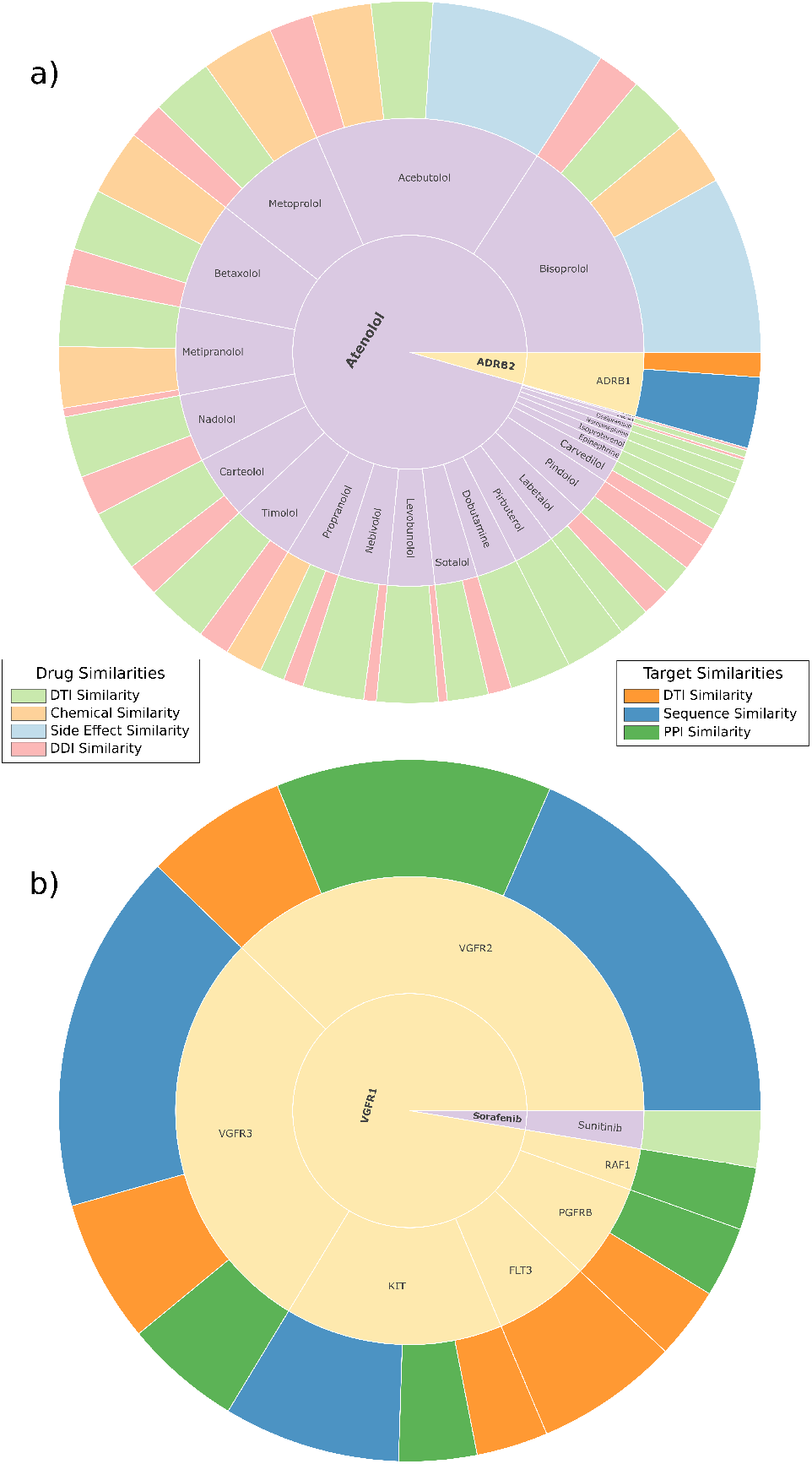
Interpretation of Inception’s predicted scores. The final score for a DTI can first be interpreted in terms of the drug and target self-representation models. These contributions are represented by the inner-most levels in the sunburst plots. The next level shows the aggregated similarities of each drug, for the drug self-representation model, and of each target, for the target self-representation model. The last and outer-most level shows the breakdown of each type of side information used to calculate the similarities. (**a**) Sunburst plot of the Atenolol-ADRB2 DTI, the top- 10 prediction for our prospective evaluation set. 95% of the score is explained by the drug self-representation model and 5% by the target self-representation model. For the drug’s piece we observe the total similarity between Atenolol and the 27 drugs that target ADRB2 in 2011. Similarly, for the target’s side we observe the similarity between ADRB2 and the only known target of Atenolol in 2011. Each drug or target can then be further analysed to quantify the relative importance of each side information. (**b**) Sunburst plot of the Sorafenib-VGFR1 DTI, the top-10 prediction for our prospective evaluation set. For this example, the drug-self representation model only accounts for 3% of the final score and the target self-representation model accounts for the remaining 97%. In the breakdown of the drug’s piece we observe the breakdown of the similarity between Sorafenib and the only drug that targets VGFR1 in the 2011 dataset. In the target’s side we observe the breakdown of the similarities between VGFR1 and the other known targets of Sorafenib in 2011.

Inception has the ability to further inform us about the additive contribution of each type of similarity information in the drug and target self-representation models. The sunburst plots in Figure 4 (a) show a breakdown of the score provided by each model. The drug self-representation model predicts the Atenolol-ADRB2 association by linearly combining similarities between Atenolol and other drugs that are known to target ADRB2 (see Equation 4). We observe that Bisoprolol and Acebutolol are responsible for 33% of the drug self-representation score. Interestingly, the contributions of Bisoprolol and Acebutolol are mainly explained by their similarities to the side effect profiles of Atenolol. These drugs, in fact, also belong to the family of beta-blockers, and are known to have similar side effect profiles. However, notice that the contribution of other beta-blocker drugs, and other types of information, are also important for the drug self-representation score.

Although the target self-representation model only contributes to 5% of the final predicted score for Atenolol-ADRB2, it can still provide useful biological insight. The target self-representation model predicts the Atenolol-ADRB2 association by linearly combining similarities between ADRB2 and other targets of Atenolol. Since Atenolol has only one known target (ADRB1), the target self-representation score is mainly explained by the similarity in protein sequence between ADRB1 and ADRB2. These proteins, in fact, participate in the same biological process of mediating the catecholamine-induced activation of adenylate cyclase through the action of G proteins.

To further illustrate the interpretation of the scores provided by Inception, we select a different DTI from our prospective evaluation set. The new association is the drug Sorafenib (DB00398) with the vascular endothelial growth factor receptor 1 (VGFR1). This DTI is the top-10 prediction of Inception with a score of 0.583. In this example, 97% of the score is explained by the target self-representation model and only 3% by the drug self-representation model (see Figure 4b).

The sunburst plot of Figure 4 (b) shows the breakdown of the score provided by each self-representation model. The target self-representation model predicts the DTI by linearly combining similarities between VGFR1 and the known targets of Sorafenib (see Equation 4). We observe that two targets stand out from the rest: VGFR2 and VGFR3. The contributions of these two targets are primarily explained by their protein sequence similarity to VGFR1. These proteins, in fact, are known to play major roles in the same biological processes and share similar molecular functions.

Notice how in the previous example, the final score was mostly explained by the drug self-representation model, in this case it only contributes 3% of the final predicted score for Sorafenib-VGFR1. However, it can still provide useful biological information about the prediction. The drug self-representation model predicts the DTI by linearly combining similarities between Sorafenib and other drugs that are known to target VGFR1. Since VGFR1 was only targeted by one drug, the drug self-representation score is explained fully by the DTI similarity between Sorafenib and Sunitinib. It is interesting to notice that these two drugs are both approved for the treatment of renal cell carcinoma. These two drugs actually have the majority of their targets in common, including VGFR2 and VGFR3. This fact is adding some extra weight to the explanation of the final predicted score for the Sorafenib-VGF1 DTI. Thus, by analysing the breakdown of the scores, we can conclude that our model is predicting the new DTIs by integrating evidence from heterogeneous types of biological evidence.

## 5 Discussion

The elucidation of drug targets remain central in mechanism-based drug discovery [Santos et al., 2017]. Finding drug targets have been compared to finding a needle in a haystack. Yet, in systematically finding drug targets lies the answer to finding treatments for many human diseases.

Here we present Inception, an interpretable machine learning model for predicting missing drug-target interactions. Inception outperforms recent state-of-the-art machine learning approaches by a wide margin. Machine learning approaches, such as DTINet and neoDTI, can provide black-box predictions but scientists are left with the difficult task of understanding them. We developed Inception to provide straightforward interpretation of DTI predictions on the basis of known biological information.

A model that attempts to effectively predict missing drug-target interactions needs to integrate available information about drugs and its protein targets. However, the integration of multiple heterogeneous information into a machine learning model is not straightforward. DTINet [Luo et al., 2017] and NeoDTI [Wan et al., 2019] have successfully integrated heterogeneous information into powerful matrix decomposition and deep learning models but at the expense of interpretability. Instead, Inception is capable of providing interpretability without compromising prediction accuracy; a common misconception in the literature [Rudin, 2019].

Often, integrative machine learning models cannot be applied when the side information about a drug or a target is not available; thus limiting their real applicability in certain cases. However, Inception can still be applied in the cases where a drug (or target) have a small number of associations, but not other side information is available. We verified that the performance of Inception remains robust even when using only the matrix **X** alone for the predictions, or when adding one side information at a time (see Supplementary Figure 1).

An innovative aspect of our self-expressive model is that it learns similarities between drugs and between protein targets by a weighted linear combination of heterogeneous information. Previous models based on self-expressiveness [Fan and Chow, 2017, Frasca et al., 2019], do not effectively integrate similarity matrices in the learned self-representation matrices. Our algorithm in Equation 3 is easy to implement, runs in a few seconds, and do not require complicated optimisation algorithms. We envisage its use in other important problems where the task is to predict missing associations between pairs of objects. This is, for instance, in the problems of drug-side effect prediction [Galeano et al., 2020], drug repositioning [Frasca et al., 2019], and disease-gene prediction [Cáceres and Paccanaro, 2019].

## Supplementary Material

### Supplementary Figures

**Figure 5:**
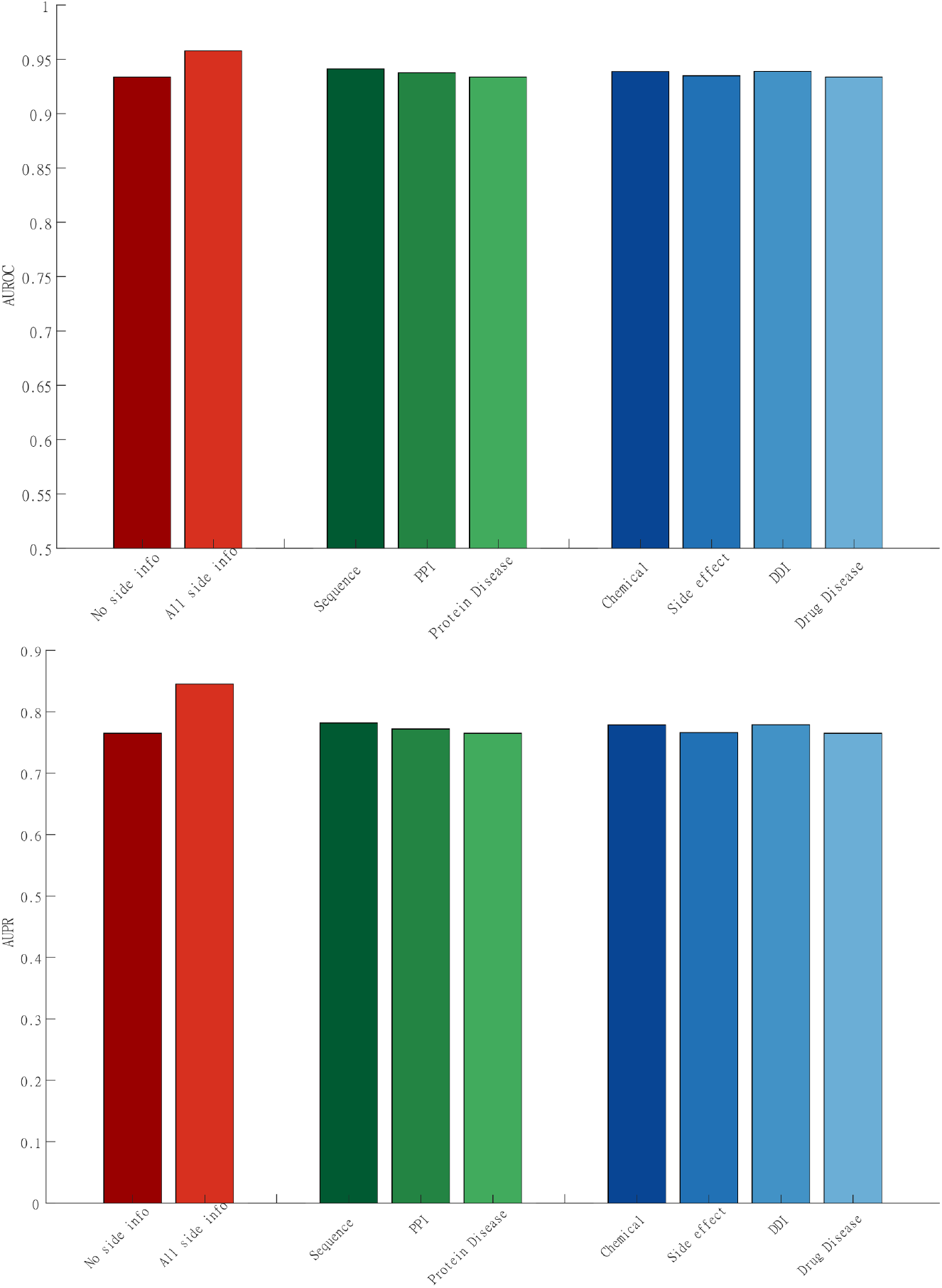
Performance of Inception in a ten-fold cross-validation procedure. The binary classification performance is shown when training the model using only one protein side information at a time (shades of green), when using one drug side information at a time (shades of blue) and using all the available side information (bright red). (**Top**) mean AUROC; (**Bottom**) mean AUPR.

### Supplementary Tables

**Table 1:** Datasets.

| Matrix | Rows | Columns | Number of known associations |
| --- | --- | --- | --- |
| Drug-target interaction | 549 | 424 | 1,923 |
| Chemical similarity | 549 | 549 | 301,401 |
| Drug Side effect | 549 | 3588 | 65,240 |
| Drug-drug interaction | 549 | 549 | 6,078 |
| Drug-disease association | 549 | 5022 | 170,188 |
| Sequence similarity | 424 | 424 | 179,776 |
| Protein-Protein Interaction | 424 | 424 | 1,029 |
| Protein-Disease interaction | 424 | 5022 | 476,457 |

**Table 2:**
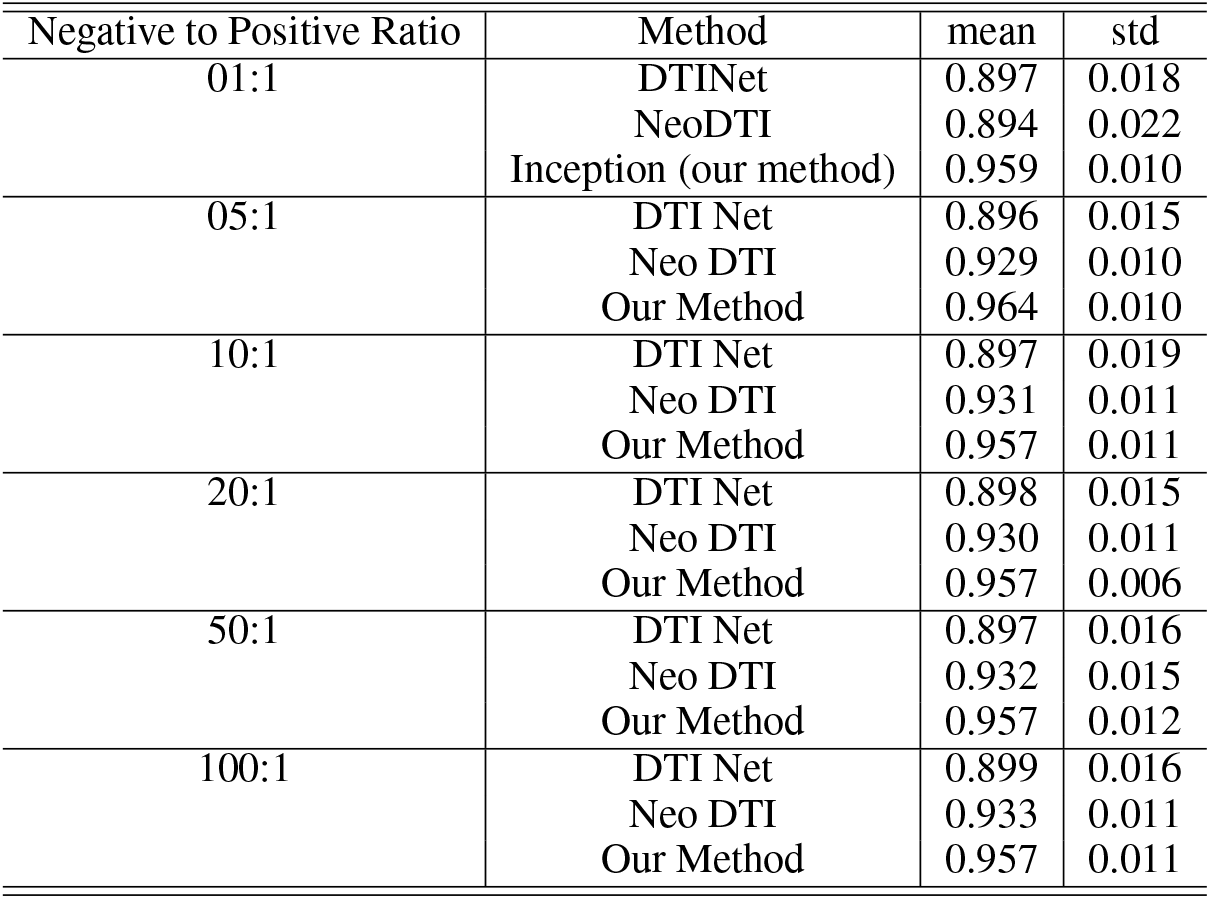
AUROC at different negative to positive ratios.

**Table 3:** AUPR at different negative to positive ratios.

| Negative to Positive Ratio | Method | mean | std |
| --- | --- | --- | --- |
| 01:1 | DTI Net | 0.915 | 0.012 |
|  | Neo DTI | 0.901 | 0.019 |
|  | Our Method | 0.962 | 0.007 |
| 05:1 | DTI Net | 0.789 | 0.026 |
|  | Neo DTI | 0.839 | 0.021 |
|  | Our Method | 0.900 | 0.019 |
| 10:1 | DTI Net | 0.718 | 0.034 |
|  | Neo DTI | 0.795 | 0.025 |
|  | Our Method | 0.845 | 0.019 |
| 20:1 | DTI Net | 0.643 | 0.019 |
|  | Neo DTI | 0.755 | 0.026 |
|  | Our Method | 0.796 | 0.022 |
| 50:1 | DTI Net | 0.530 | 0.037 |
|  | Neo DTI | 0.695 | 0.040 |
|  | Our Method | 0.716 | 0.039 |
| 100:1 | DTI Net | 0.440 | 0.031 |
|  | Neo DTI | 0.642 | 0.033 |
|  | Our Method | 0.653 | 0.043 |

**Table 4:** AUROC when removing different similarities.

| Removed Association | Method | mean | std |
| --- | --- | --- | --- |
| Chem | DTINet | 0.826 | 0.042 |
|  | IncEpTion | 0.936 | 0.027 |
|  | NeoDTI | 0.887 | 0.018 |
| ChemProt | DTINet | 0.742 | 0.078 |
|  | IncEpTion | 0.906 | 0.045 |
|  | NeoDTI | 0.789 | 0.049 |
| DDI | DTINet | 0.855 | 0.041 |
|  | IncEpTion | 0.940 | 0.020 |
|  | NeoDTI | 0.895 | 0.030 |
| Prot | DTINet | 0.793 | 0.061 |
|  | IncEpTion | 0.918 | 0.032 |
|  | NeoDTI | 0.846 | 0.032 |
| SideEffect | DTINet | 0.895 | 0.021 |
|  | IncEpTion | 0.956 | 0.013 |
|  | NeoDTI | 0.930 | 0.013 |

**Table 5:** AUPR when removing different similarities.

| Removed Association | Method | mean | std |
| --- | --- | --- | --- |
| Chem | DTINet | 0.554 | 0.070 |
|  | IncEpTion | 0.759 | 0.067 |
|  | NeoDTI | 0.688 | 0.064 |
| ChemProt | DTINet | 0.346 | 0.121 |
|  | IncEpTion | 0.596 | 0.100 |
|  | NeoDTI | 0.412 | 0.127 |
| DDI | DTINet | 0.591 | 0.081 |
|  | IncEpTion | 0.779 | 0.044 |
|  | NeoDTI | 0.701 | 0.055 |
| Prot | DTINet | 0.421 | 0.090 |
|  | IncEpTion | 0.635 | 0.073 |
|  | NeoDTI | 0.518 | 0.074 |
| SideEffect | DTINet | 0.711 | 0.041 |
|  | IncEpTion | 0.842 | 0.027 |
|  | NeoDTI | 0.793 | 0.029 |

